# Stochastic processes dominate community assembly in cichlid communities in Lake Tanganyika

**DOI:** 10.1101/039503

**Authors:** Thijs Janzen, Adriana Alzate, Moritz Muschick, Fons van der Plas, Rampal S. Etienne

## Abstract

The African Great Lakes are characterized by an extraordinary diversity of endemic cichlid fish species. The cause of this diversity is still largely unknown. Most studies have tried to solve this question by focusing on macro-evolutionary processes, such as speciation. However, the ecological processes determining local cichlid diversity have so far been understudied, even though knowledge on these might be crucial for understanding larger scale biodiversity patterns.

Using trait, environmental and abundance data of cichlid fishes along 36 transects, we have studied how differences in local environmental conditions influence cichlid community assembly in the littoral of Lake Tanganyika, Zambia. We investigated changes in average trait values and in trait-based community assembly processes along three key environmental gradients.

Species diversity and local abundance decreased with increasing sand cover and diet-associated traits changed with depth. Analyses on within-community trait diversity patterns indicated that cichlid community assembly was mainly driven by stochastic processes, to a smaller extent by processes that limit the similarity among co-existing species and least by filtering processes that limit the range of species traits occurring in an environment. Despite, the low impact of habitat filtering processes, we find community dissimilarity to increase with increasing environmental difference.

Our results suggest that local environmental conditions determine cichlid abundance, while the predominance of stochastic community assembly across all environments explains why the communities with the highest abundances contain most species.

## INTRODUCTION

The stunning diversity of cichlid fish in the African Rift lakes has fascinated scientists during the past 50 years (Brooks 1950; Fryer & Iles 1972; Coulter 1991; Kocher 2004; Wagner *et al.* 2012). In contrast to the multitude of research focusing on the evolutionary explanations of cichlid diversity (Sturmbauer *et al.* 2001; Salzburger *et al.* 2005; Muschick *et al.* 2012; Brawand *et al.* 2014), there are fewer studies aiming at understanding ecological mechanisms underlying local coexistence and diversity of cichlid species. Empirical studies on local scale diversity have focused either on temporal trends (Hori *et al.* 1993; Takeuchi *et al.* 2010), the impact of human disturbance (Alin *et al.* 1999), opportunities for relieving fishing efforts (Duponchelle *et al.* 2003; Weyl *et al.* 2005), the impact of protected areas on cichlid communities (Sweke *et al.* 2013) or have been restricted to descriptions only (Hori *et al.* 1983; Kuwamura 1987; Van Steenberge *et al.* 2011). While these studies have shown interesting patterns of local cichlid communities, no studies so far have attempted to understand the processes underlying local community composition patterns.

Community assembly is generally viewed from two alternative perspectives, either from a niche-based perspective, or a neutral perspective. The niche-based hypothesis postulates that species are adapted to their local environment and occupy a species’ specific niche: a set of conditions in which the species thrives, and outcompetes other species (Hutchinson 1959; Tilman 1982; Chesson 2000). The traits of a species reflect the adaptation of a species to its own niche, and studying patterns in trait-distributions can inform us about any underlying processes driving species-coexistence and community composition. In benign environments that do not pose strong requirements on traits, the niche-based hypothesis predicts that species are abundant, and that the presence or absence of species’ traits is mainly shaped through interactions with each other, rather than interactions with the abiotic environment. Due to competitive exclusion of species with overlapping niches or due to exclusion of species with shared specialist predators, niche-based assembly is expected to lead to an *increase* in trait diversity of co-occurring species in benign environments (Macarthur & Levins 1967; Mayfield & Levine 2010). In harsh environments, species with traits that make them intolerant to stress, herbivory and/or predation pressures might be excluded from a local community, leading to a *reduced* observed trait diversity of co-occurring species (Weiher & Keddy 1995; Cornwell *et al.* 2006).

In contrast, the neutral hypothesis, which treats all individuals from all species equally, explains community composition from a stochastic point of view, where local abundance of a species is the outcome of stochastic birth, death and migration over time (Hubbell 2001; Rosindell *et al.* 2011, 2012). Local community composition is assumed to be a dynamic equilibrium between random immigration from the species pool, and local ecological drift. The neutral hypothesis acknowledges that there might be benign and harsh environments, but that these environments affect all individuals equally. As a consequence, benign environments have more individuals than stressful environments, but both benign and stressful areas contain individuals that form a (dispersal-limited) random subset of the species pool. The null expectation is then that areas with high abundances also have high diversity, as a result of random sampling.

Here we investigate how niche-based and neutral-based community assembly shapes biodiversity in cichlid fishes across three environmental gradients: depth, sand cover and topographical complexity. If community assembly is highly niche-driven, we firstly expect that decreasing algae densities with depth provide less scope for algae-feeding cichlids and that hence, communities in deep areas are predominantly assembled by filtering processes. Secondly, because many cichlids are highly dependent on rocks for either food (as substrate for algae growth) or for shelter (Konings 2005), we expect that an increase in sand cover, which brings a reduction of available rock surface, will increase habitat filtering and decrease diversity. However, males of several cichlid species use sand to construct bowers to attract females and are thus highly dependent on sand, which could counteract filtering processes and possibly drive limiting similarity processes (Sefc 2011). Thirdly, we expect that areas with high topographical complexity induce higher levels of limiting similarity and higher diversity, because a more complex habitat provides more shelter and a larger surface area for algae growth. Furthermore, complex habitats are often associated with a reduction in territoriality (Danley 2011), a larger number of niches (Willis *et al.* 2005) and higher diversity (Ding *et al.* 2014).

Alternatively, if neutral-based processes dominate, we expect resource-rich areas, such as shallow, rocky and complex habitats that allow for abundant algae growth, to show highest diversity, as these habitats can sustain the highest number of individuals. Furthermore, we expect these habitats to have the highest species diversity, and we expect that the trait compositions of such communities to reflect a random subset of the larger species pool.

## METHODS

### Abundance data

Abundance and composition data of cichlids was collected in Lake Tanganyika, near Kalambo Lodge (8°37′22.29"S, 31°12′1.89"E), Zambia, Africa (Figure 1) using SCUBA diving. In total, 12 different sites were sampled, with a total of 36 transects, which were placed parallel to the shore (Figure 1, Table 2). Individuals were recorded along 20m x 4m transects by two divers in two steps: first, all individuals within 2 meters on one side of the transect were sampled. After an interval of 10 minutes, all individuals within 2 meters on the other side of the transect were sampled.

**Figure 1.**
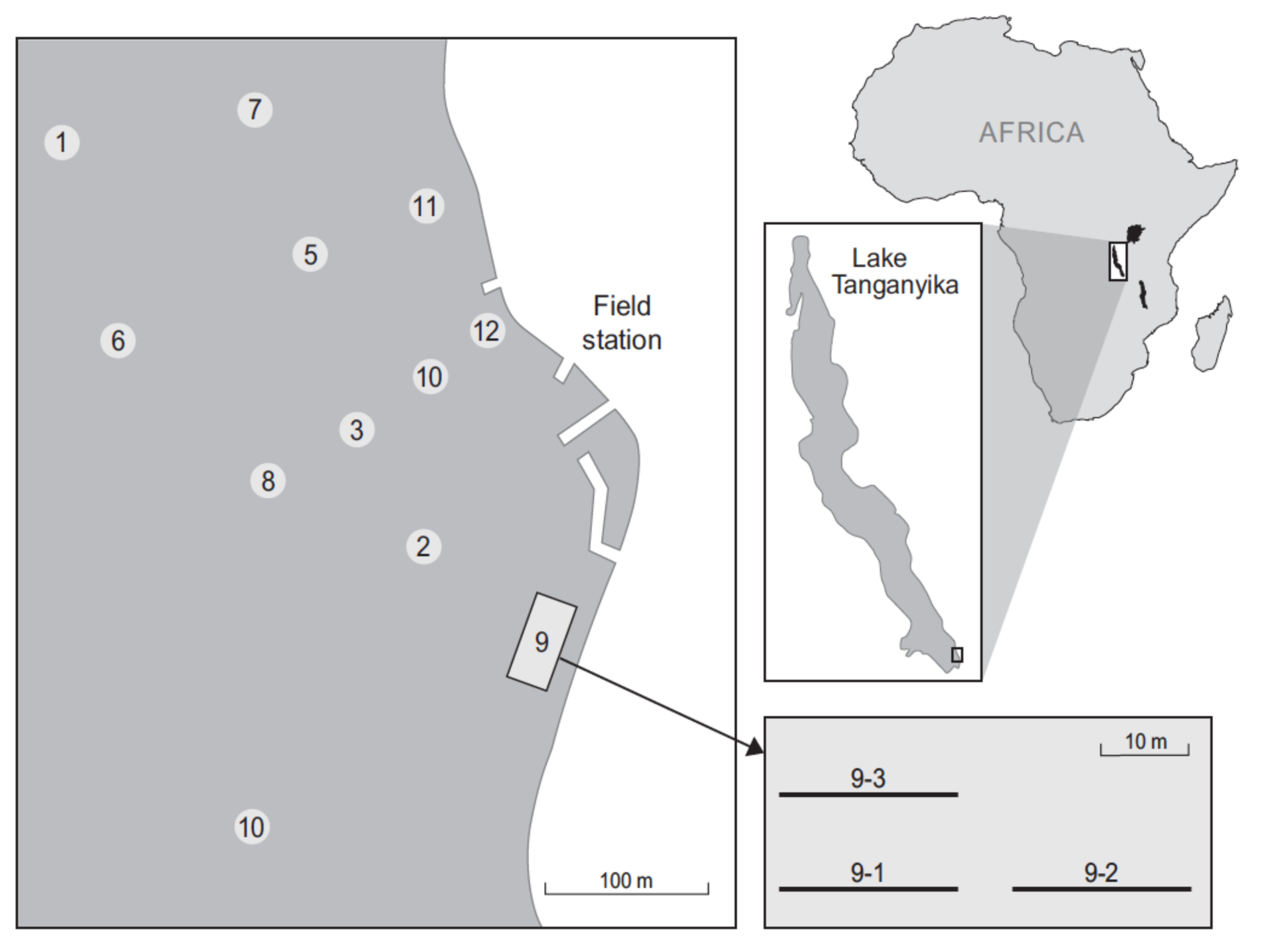
Sketch of the sampling positions in front of the Kalambo Lodge, located in the south of Lake Tanganyika. Relative position of the transects at every site is indicated in the left panel. Numbers in the figure refer to site numbers in Tables S1, S2 and S3.

### Environmental data

At each transect, we measured three different ecological variables. Firstly, we recorded the depth at the beginning and the end of each transect. Secondly, we took 40 photos of 50 x 50 cm quadrats per transect, to estimate the percentage of sand cover (20 photographs on the left side of the transect line, and 20 photographs on the right side). Percentage of sand cover per quadrat was calculated using an image analysis script in MATLAB (supplementary information). The average proportion of sand across all photographs taken along the transect was used to quantify sand cover of the transect. Thirdly, we recorded the topographical complexity of the substrate using a variation of the ‘chain method’ (Risk 1972). Along the transect, a link-brass chain was laid over the substrate, such that the chain would closely follow the contour of the substrate (Shumway *et al.* 2007). Topographical complexity was calculated as the ratio between the distance following the contour (measured with the chain) and the horizontal linear distance. High values indicate high relief/complexity, caused by an alternation between large rocks and small rocks or sand, while low values indicate low relief/complexity.

### Species traits

Data on mean trait values of each cichlid species as reported in Muschick et al. (2012, 2014) were collated for 11 traits: standard length, total length, weight, stable isotope ratios of carbon and nitrogen, lower pharyngeal jaw height, lower pharyngeal jaw width, gut length, lower pharyngeal jaw shape and body shape. In order to obtain a set of traits that does not strongly correlate with each other, to avoid the overemphasis of the importance of some traits over others, we only used a subset of these traits. Firstly we checked the Pearson Rank correlation between traits and removed one of the traits that correlated strongly. Total length and weight were found to correlate strongly with standard length (standard length vs total length: R^2^ = 0.98, standard length vs weight: R^2^ = 0.75), and we decided to only use standard length. Furthermore, lower pharyngeal jaw (LPJ) width and lower pharyngeal jaw height were found to correlate with each other (R^2^ = 0.67) and to correlate with standard length (lpj height vs standard length: R^2^ = 0.44, lpj width vs standard length: R^2^ = 0.37). Partly to avoid overemphasis of pharyngeal jaw traits, and partly because of the high correlation coefficients we decided to omit both these traits. Lower pharyngeal jaw shape and body shape were assessed using landmark-based geometric morphometric methods. *xy* coordinates of the landmarks were combined into PCA components for both body shape and lower pharyngeal jaw shape. From these PCA components we included the first PCA component for both body and lower pharyngeal jaw shape, which contained 37% and 49% of the variation for body and pharyngeal jaw shape respectively. The inclusion of more PCA components would allow us to cover a larger part of variance in shapes, but would overemphasize the importance of shape relative to the other traits.

The final trait set thus consisted of 6 traits: standard length, stable isotope ratios of carbon and nitrogen, gut length, the first PCA component of LPJ shape and the first PCA component of body shape. Missing trait values were imputed using the MICE package for R (Buuren & Groothuis-Oudshoorn 2011). The MICE imputation process uses a Gibbs sampler technique to impute the missing data, assuming a multivariate distribution for the missing data.

### Trait-based community assembly

In order to infer the relative contribution of limiting similarity, habitat filtering and dispersal assembly we used the STEPCAM approach (Van der Plas *et al.* 2015). The STEPCAM model is a STEPwise Community Assembly Model, which implements community assembly by applying three types of processes that mediate species selection from the species pool to the local community: ’filtering processes’, ’limiting similarity processes’ and ’stochastic processes’. Starting with all observed species in the dataset, species are removed in a stepwise fashion until the number of species observed in the local community is reached. Removal of species occurs either 1) because their traits are too dissimilar from the observed mean trait distribution in the community, which is assumed to be the habitat optimum (‘filtering’), 2) because their traits are too similar to the other remaining species (‘limiting similarity’), or 3) due to a stochastic event, which results in a random removal step, where the probability of removal is proportional to the number of local communities in the dataset where the species is observed, which is used as a proxy for the species pool. The STEPCAM model was fitted using Approximate Bayesian Computation, where using the model, data is simulated and compared with the observed data. Comparison between simulated and observed data occurred through comparing four summary statistics, Functional Richness, Functional evenness, Functional divergence and community trait means (Villéger et al. 2008, van der Plas et al. 2015). We applied a Sequential Monte Carlo algorithm (ABC-SMC) using the function STEPCAM_ABC from the package STEPCAM (Janzen & van der Plas 2014). We used 1000 particles and a final acceptance rate of 1 in 20,000. The reported estimates for dispersal assembly, filtering and limiting similarity are mean estimates over 3 replicate STEPCAM runs, with the random number generator seeded with different seeds for each replicate.

### The effect of the environment on community trait means

In order to understand changes in trait values that might be linked to changes in the environment, we analyzed the relationship between the different traits and environmental variables. We calculated, per transect, community-level weighted trait means (CWM), where trait values are weighted by the relative frequency of the species in the transect (Lavorel *et al.* 2008). CWM values were calculated using the function ‘dbFD’ from the R package FD (Laliberté *et al.* 2014). We used Linear Mixed Models to test how CWM values changed over environmental gradients. We constructed full models, where CWM values were treated as dependent variable, the three habitat components as fixed effect, and the site as a random effect. We then used stepwise removal to remove non-significant effects. Conditional R^2^ values were calculated following Nakagawa and Schielzeth (2013).

We repeated this approach using unweighted trait means, thus treating every species equally and disregarding differences in abundance. Using unweighted trait means has two benefits over using weighted means: firstly it does not overemphasize species that occur in a clustered fashion, such as the breeding complexes of *Neolamprologus pulcher*. Secondly it does not underestimate species that, although occurring in low abundance, influence species around them strongly, for instance due to territoriality or aggressive behavior, such as *Plecodus straeleni* (Boileau *et al.* 2015).

### Community dissimilarity

To assess the simultaneous effect of all three environmental gradients on community composition, we compared community dissimilarity between all communities. We constructed an environmental distance score by calculating the distance between all transects for each gradient (depth, sand and complexity). To ensure that all environmental factors had a similar weight on environmental heterogeneity, we normalized the distance scores by the maximum distance, such that all individual distance scores where between -1 and 1. We then obtained the total normalized environmental distance by calculating the Euclidian distance by taking the square root of the sum of squared distance scores.

## RESULTS

### Species compositions

We recorded on average 22 species per transect, and an accumulated total number of 53 species (Table S3). *Telmatochromis temporalis* was the most common species, contributing 12% of 4926 recorded individuals. The seven most common species combined accounted for 50% of all observed individuals (*Telmatochromis temporalis, Variabilichromis moorii, Tropheus moorii, Neolamprologus pulcher, Interochromis loocki, Telmatochromis vittatus* and *Xenotilapia boulengeri*), whilst the 24 most common species accounted for 90% of all individuals.

Transects with a higher sand cover had a lower number of individuals (*R*^*2*^ = 0.41, p = 0.021), and a lower species richness (*R*^*2*^ = 0.47, p = 0.001) (Table 1, Figure 2). Neither depth nor habitat complexity had significant effects on abundance or species richness (Table 1, Figure 2).

**Table 1.** Significant components of linear mixed-effects models, where species traits were used as response variable, the habitat components as predictor variables, and the sampling site as a random effect. Only those components that were significant after stepwise removal of all non-significant components are reported. The conditional *R*^2^ of the final model is reported in the last column. Used abbreviations: LPJ = Lower Pharyngeal Jaw, PCA = Principal Component Axis.

| Response variable | Depth | Sand | Complexity | $R^2$ |
| --- | --- | --- | --- | --- |
| Species richness |  | -0.1143 |  | 0.47 |
| <sup>10</sup>log abundance |  | -0.0033 |  | 0.67 |
| **Weighted Trait Means** |  |  |  |  |
| Standard Length |  | 0.1386 |  | 0.54 |
| Total Length |  | 0.1652 |  | 0.57 |
| Weight |  | 0.0505 | 5.1724 | 0.40 |
| δ15N | 0.0553 |  | -0.3885 | 0.78 |
| δ13C | -0.1754 |  |  | 0.73 |
| LPJ height |  | 0.0074 |  | 0.69 |
| LPJ width | 0.0002 | 0.0001 |  | 0.72 |
| Gut Length |  |  | 171.79 | 0.52 |
| LPJ PCA axis 1 | -0.2727 | -0.0185 |  | 0.84 |
| LPJ PCA axis 2 | -0.0611 |  | 2.3922 | 0.54 |
| LPJ PCA axis 3 |  | -0.0100 | 0.8428 | 0.56 |
| Body PCA axis 1 |  | 0.0404 |  | 0.67 |
| Body PCA axis 2 | 0.0770 | -0.0092 |  | 0.67 |
| Body PCA axis 3 | -0.0989 | 0.0141 | -1.6877 | 0.86 |
| **Unweighted Trait Means** |  |  |  |  |
| Standard Length | -0.2700 | 0.0413 | 7.7057 | 0.54 |
| Total Length |  |  | 7.3057 | 0.48 |
| Body Weight | -0.3291 |  | 5.3659 | 0.58 |
| δ15N | 0.0366 |  |  | 0.76 |
| δ13C | -0.0899 | -0.0066 |  | 0.78 |
| LPJ height | -0.0109 |  |  | 0.32 |
| LPJ width | -0.0002 |  |  | 0.29 |
| Gut Length | -8.8453 |  | 130.8656 | 0.58 |
| LPJ PCA 1 | -0.0840 | -0.0109 | -1.3614 | 0.78 |
| LPJ PCA 2 | -0.0880 |  | 0.8963 | 0.54 |
| LPJ PCA 3 |  | -0.0039 | 0.7303 | 0.51 |
| Body PCA 1 | 0.0668 | 0.0126 |  | 0.65 |
| Body PCA 2 | 0.0667 | -0.0041 |  | 0.67 |
| Body PCA 3 | -1.5261 |  |  | 0.60 |

**Figure 2.**
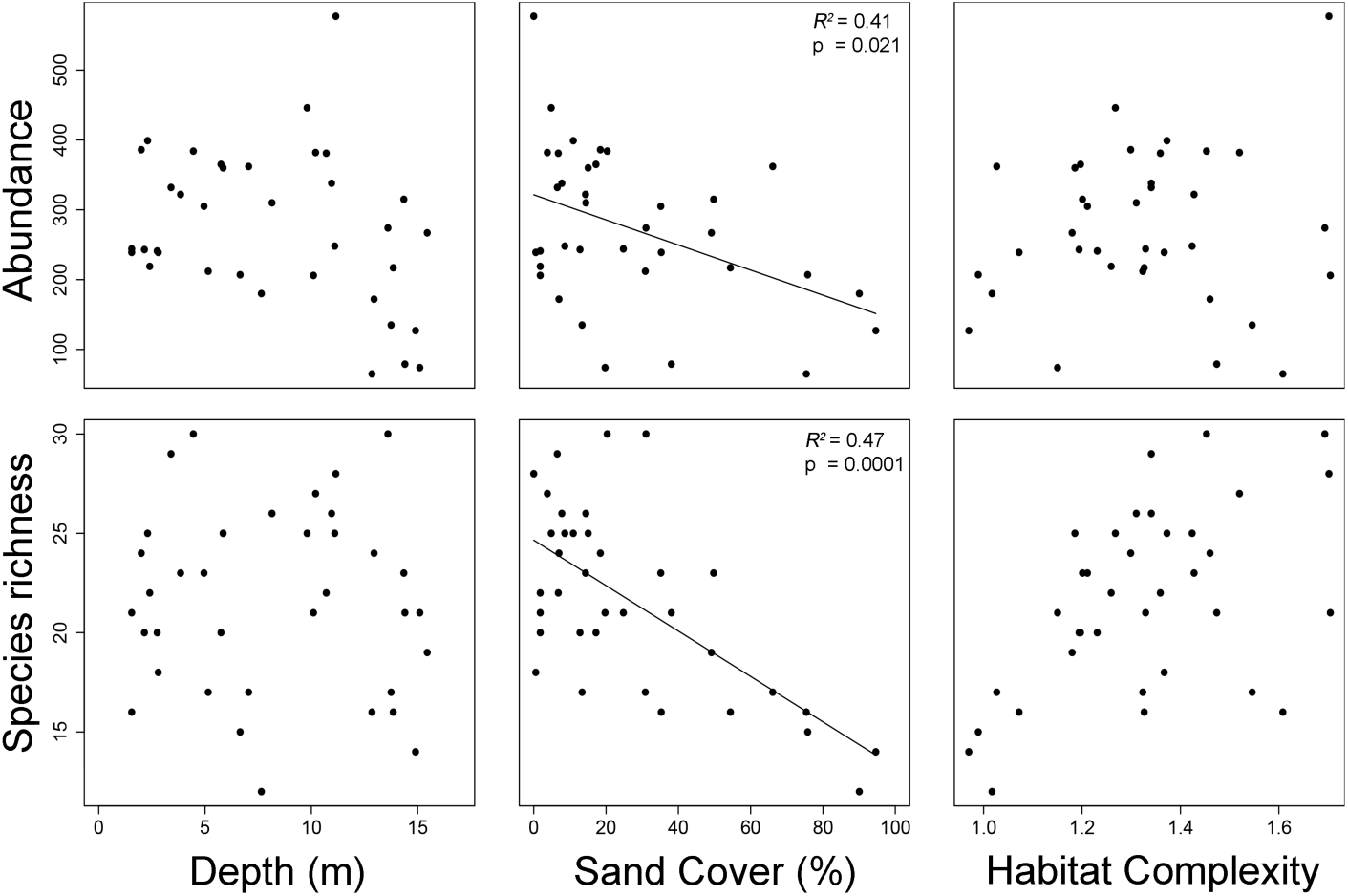
Changes in abundance and in species richness across the three habitat gradients. Points depict the different transects. Significant correlations (table 1) are plotted as a line.

We compared the Bray-Curtis dissimilarity between transects with the normalized Euclidian distance between the environmental conditions at those transects. Bray-Curtis dissimilarity was found to correlate significantly with the normalized environmental distance score (figure 3, *R*^*2*^ = 0.59, Mantel test, mantel-r statistic = 0.766, p = 0.00001, 100.000 permutations).

**Figure 3.**
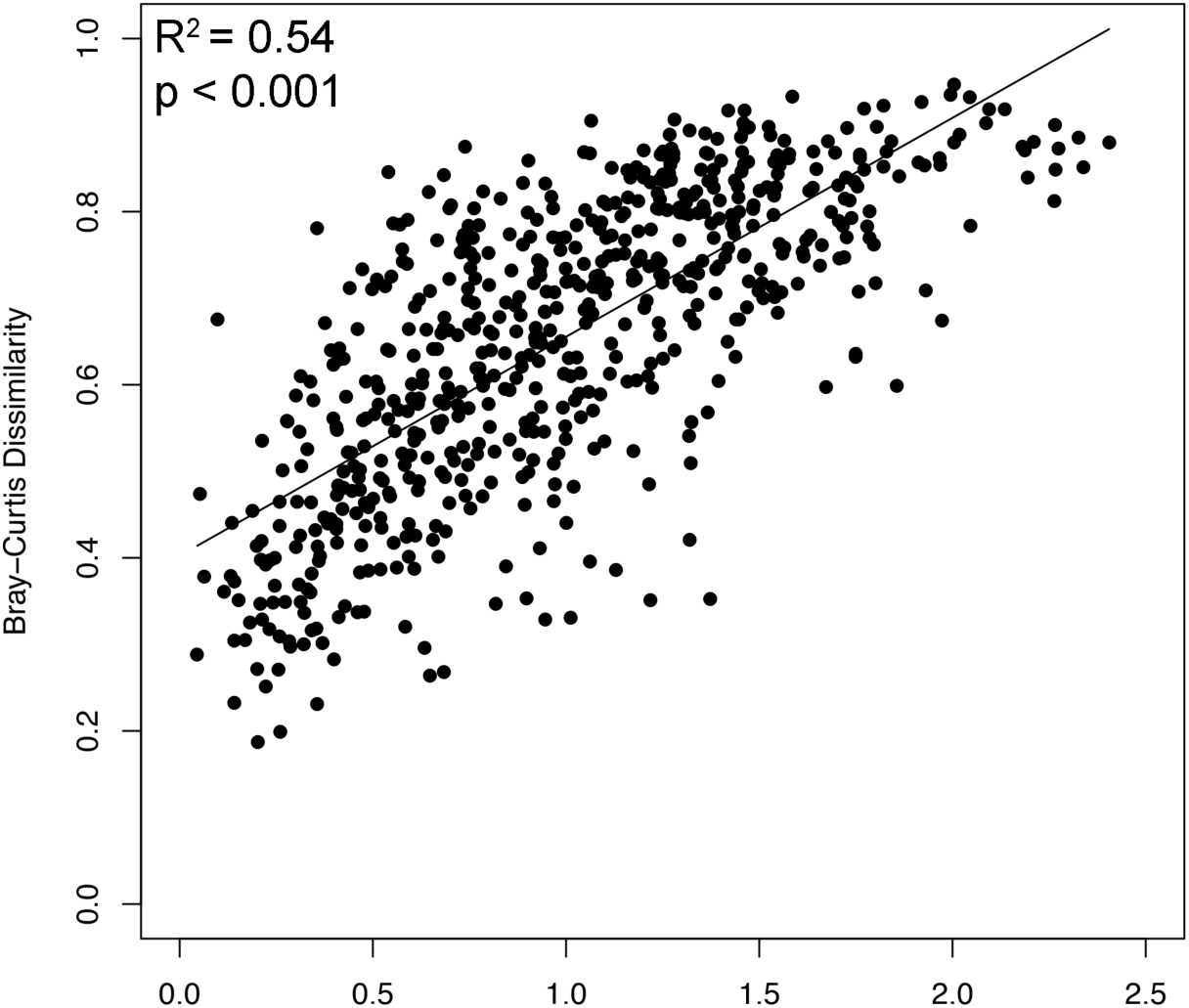
Changes in Bray-Curtis dissimilarity between sampled transects versus the normalized environmental distance between sampled transects. Normalized environmental distance is the Euclidian distance between two plots, where the environmental distances are normalized by the maximum value recorded across all plots.

### Relative contribution of community assembly processes

Fitting STEPCAM to the trait distributions of the 36 different transects yielded an average contribution of dispersal assembly steps of 72%, an average contribution of habitat filtering steps of 9% of the steps and 19% of removal steps due to limiting similarity (Figure 4). We found no significant correlations between relative contribution of either of the three processes and any of the three habitat gradients (Figure 5).

**Figure 4.**
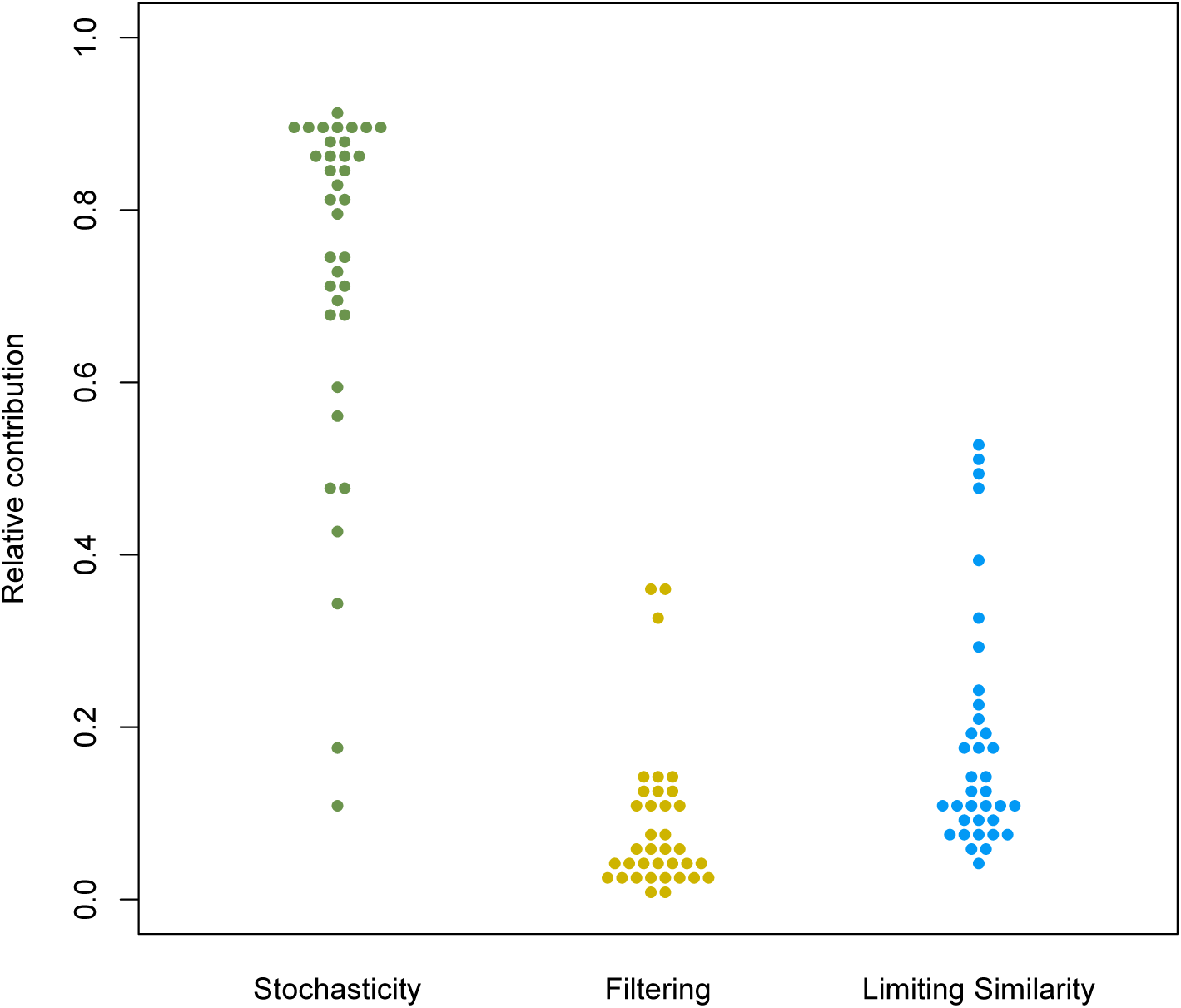
Relative contribution of stochasticity, habitat filtering and limiting similarity steps across all 36 sites.

**Figure 5.**
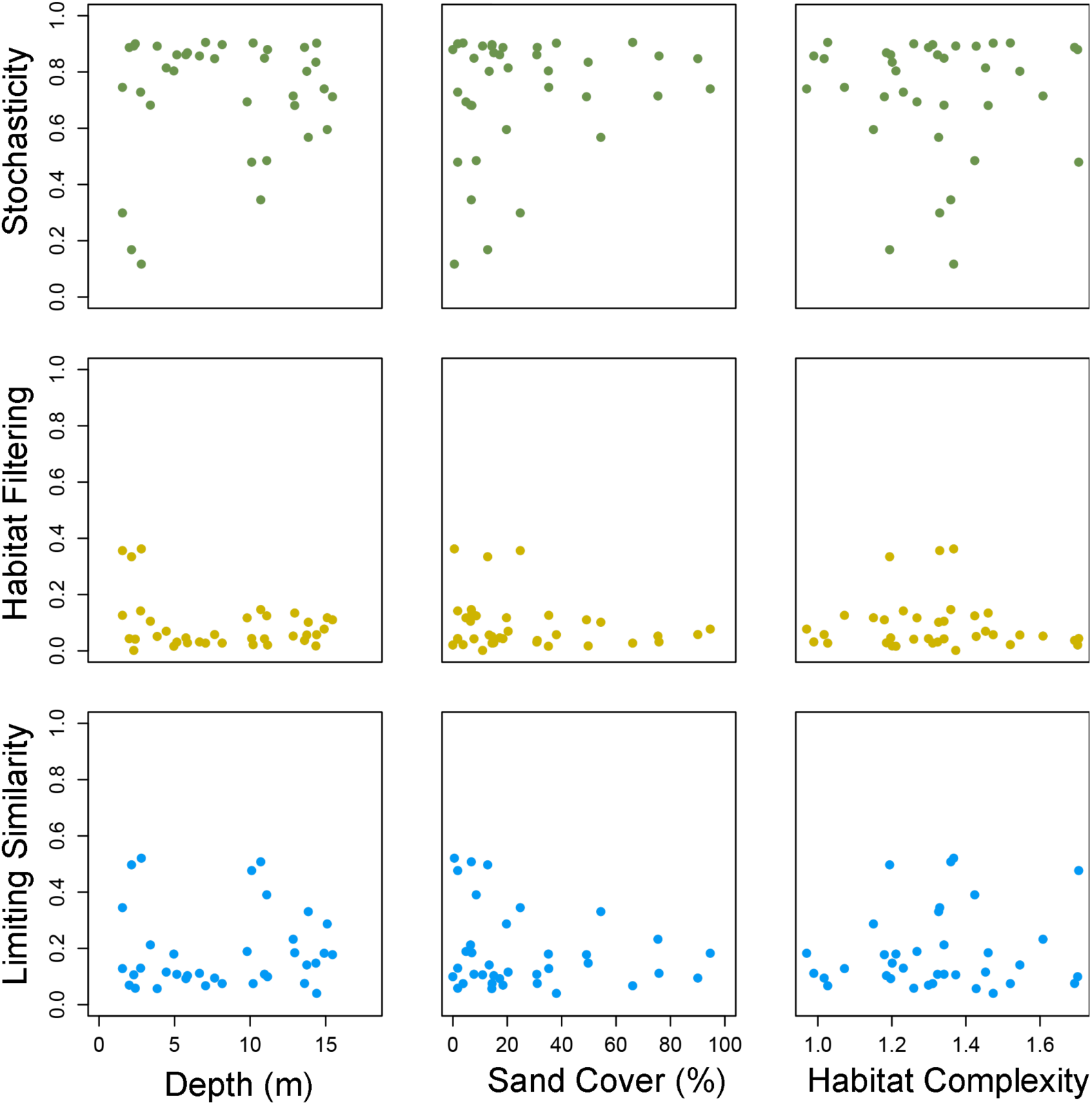
Relative contribution of stochasticity, habitat filtering and limiting similarity steps as estimated using STEPCAM, plotted against the three measured habitat components: depth, sand cover and complexity. None of these relationships was significant.

### Relationships between traits and habitat components

We found that the three habitat gradients explained only a low proportion of variation (*R*^2^ around 0.5) of community-weighted means (CWM) of cichlid traits (Table 1). We observed high *R*^*2*^ values (*R*^*2*^>0.7) for d15N content, d13C content, lower pharyngeal jaw width, the first axis of the PCA of the Lower Pharyngeal Jaw shape, and the third axis of the PCA of the body shape (Table 1). Using community unweighted trait means we find similar effects, with similar *R*^2^ scores (Table 1), suggesting that weighting the trait values by abundance does not influence our findings.

## DISCUSSION

We have investigated whether community assembly in cichlid communities at the littoral of Lake Tanganyika, Zambia is mostly driven by niche-based processes, or by neutral-based processes. In line with the neutral hypothesis, we found that while abundance and richness of cichlids became higher in benign conditions, within-community traits distributions were highly similar to random samples from the species pool distribution. Despite some modest relationships between average trait values and habitat components, the within-community trait distribution reflects a dominance of stochastic processes in community assembly. Niche-based processes only contributed minimally to community assembly and instead, observed community patterns pointed to an important role for neutral processes in community assembly.

Although habitat filtering only contributed to community assembly to a limited extent, environmental conditions were not without influence on local community patterns. We found that an increase in sand cover was correlated with a lower total abundance in the local community. This could be an effect of a reduction in food resources, as there is less substrate for algae growth (a common food source for many cichlids). However, individuals of most species showed similar trends, which explains why a) both in areas with high and low sand cover local communities resembled a random subset of the species pool, and b) diversity is highest in low sand areas: this is mostly the result of random sampling from the species pool. Sand cover thus drives local community patterns through neutral based processes, rather than niche-based processes.

Although we found no changes in the relative contribution of the different community assembly processes across the different environmental gradients, we did find that species compositions shifted over these gradients. With an increasing difference between the local conditions between two sites, their species composition became more dissimilar. This seems to suggest that habitat filtering does impose an effect on community composition. One explanation of our lack of finding habitat filtering effects in the STEPCAM analysis could lie in our choice of traits. In our trait-based analyses we focused on traits associated with shifts in diet and survivability (total length, gut length and pharyngeal jaw shape). We expected these traits to differ most across different habitats, as they are thought to be related to different trade-offs regarding resource retrieval and survival (Muschick *et al.* 2012; McGee *et al.* 2015). The inclusion of traits specifically linked to sexual selection, such as male coloration, sexual dichromatism or sexual dimorphism, could change our estimates and possibly reveal novel relationships between habitat components and stochasticity, habitat filtering or limiting similarity.

Even though observed community patterns were in line with a dominant role for stochastic processes in community assembly, trait-based processes were not completely absent. This was shown by our stepwise community assembly models, and confirmed by patterns in community weighted trait means over environmental gradients. We found significant trait-environment relationships for stable nitrogen and carbon isotopic signatures, lower pharyngeal jaw shape and shape of the body. The first three of these traits are associated with food uptake and diet and mainly change with depth (associated with light availability and hence algal density), thus confirming our expectation that there is a diet shift along our environmental gradients.

Here we have presented a first analysis of cichlid diversity from a community assembly perspective. We have focused on disentangling niche-based and neutral processes and found that the majority of community assembly in cichlid communities at the littoral of Lake Tanganyika, Zambia is driven by stochastic effects, rather than trait-based effects. Interestingly, the dominance of stochastic processes in shaping local diversity is in line with previous findings focusing on factors influencing cichlid diversity from a more macro-evolutionary (rather than local ecological) perspective. Using advanced regression techniques, Wagner and colleagues (2012, 2014) have shown that the success, and the diversity, of a cichlid radiation depends mainly on the depth, the size and the total received solar input of a lake. These three factors are the main drivers behind the maximum number of individuals that a lake can sustain, which in turn determines the number of species, following neutral theory (Rosindell & Cornell 2009).

Furthermore we suggest that our findings are in line with the previously hypothesized stages of adaptive radiation in cichlid fish (Kocher 2004). Kocher hypothesized that cichlids have first adapted to the local habitat (habitat filtering), after which diversification proceeded through diversification of the feeding apparatus (limiting similarity), and lastly diversification proceeded through diversification of the color pattern due to sexual selection. Our findings resonate this order, as we find that adaptations to the local habitat (habitat filtering) are currently least important in shaping diversity, followed secondly by trophic adaptations driven by competition for food (limiting similarity). Although the current contribution of community assembly processes is not necessarily a strong predictor of the evolutionary path that cichlids have followed, the resemblance is striking at least.

Here we studied cichlid biodiversity from a micro-ecological perspective, rather than from a (macro-) evolutionary perspective. Ultimately, a combination of these alternative approaches might be most effective in uncovering why the diversity of cichlids in the East African Rift lakes is so extremely high.

## ACKNOWLEDGEMENTS

TJ and AA thank Walter Salzburger and Adrian Indermaur for help during data collection. MM thanks Marius Roesti for help during data collection. Field work was supported by a grant from the Gratama Stichting, and a grant from het Schure-Beijerinck Popping Fonds. RSE and TJ thank the Netherlands Organisation for Scientific Research for support through a VIDI grant awarded to RSE.

